# Medial Olivocochlear Activity Modulates Central Auditory Synapse Structure and Function

**DOI:** 10.64898/2026.01.19.700164

**Authors:** Daniela María Chequer Charan, Fangfang Wang, María Eugenia Gómez-Casati, Haoyu Wang, Yumeng Qi, Walen Leonardo Gribaudo, Camila Catalano Di Meo, Carolina Wedemeyer, Yunfeng Hua, Ana Belén Elgoyhen, Mariano Nicolas Di Guilmi

## Abstract

In many mammals, the auditory system is immature at birth and undergoes activity-dependent refinement. The medial olivocochlear system (MOC) contributes to the maturation of central auditory connectivity by shaping spontaneous activity in the developing inner ear. Although altered MOC activity has been linked to electrical and synaptic dysfunctions in different central auditory nuclei, an in-depth analysis of the same synapse under conditions of absent versus enhanced MOC activity is lacking. In this work, we set out a physiological and structural analysis of the calyx of Held and the principal neurons of the medial nucleus of the trapezoid body (CH-MNTB) synapse at postnatal days 12-14 in three mouse models of either sex: wild-type (WT), α9 knock-in (α9KI; with enhanced MOC activity), and α9 knock-out (α9KO; which lacks MOC activity). Electrophysiological recordings in brain slices revealed a reduced synaptic strength efficacy in α9KI compared to WT, including smaller excitatory postsynaptic current (EPSC) amplitudes, stronger short-term depression during repetitive stimulation and a decrease readily releasable pool size. In contrast, α9KO mice showed minimal synaptic differences relative to WT. Serial block-face electron microscopy (SBEM) reconstructions demonstrated morphological alterations of the CH in both MOC-manipulated mouse models. However, α9KI mice exhibited the largest deviations, including fewer morphologically complex CHs and increased poly-innervation of MNTB cells. These results indicate that transient, well-regulated efferent control of cochlear activity is crucial for establishing accurate central auditory connectivity. Moreover, the enhancement of MOC activity drives more pronounced developmental changes in brainstem auditory circuitry than its absence.

**Significance Statement:** Before hearing onset, inner hair cells in altricial mammals show spontaneous electrical activity crucial for proper auditory pathway development. This activity is finely regulated by descending efferent nerves from the central nervous system during a critical developmental period, ensuring precise auditory circuit formation. Our study shows that genetically manipulating efferent function —either by eliminating it or enhancing it— leads to structural alterations and severe synaptic dysfunction within the auditory brainstem. These results indicate that transient, well-regulated efferent control of cochlear activity is crucial for establishing accurate central auditory connectivity. Importantly, enhancing efferent peripheral activity causes more pronounced central synaptic changes than its absence, highlighting the importance of balanced efferent modulation during early auditory system development for normal brainstem auditory function.

## Introduction

The critical period is a highly plastic developmental process involving maturation of brain and sensory systems. In altricial mammals, this process occurs during the first postnatal weeks via sensory-independent but activity-dependent mechanisms that reorganize immature synaptic and cellular networks (Goodman and Shatz, 1993; Hanson and Landmesser, 2004; Kirkby et al., 2013). In this regard, the auditory system is immature at birth, and spontaneous inner ear activity during the critical period drives its maturation (Kotak and Sanes, 1995; Lippe, 1995; Jones et al., 2007; Sonntag et al., 2009; Tritsch et al., 2010). In the pre-hearing neonatal cochlea, ATP released from supporting cells triggers calcium action potentials in inner hair cells (IHC), generating sound independent spontaneous activity (Kros et al., 1998; Glowatzki and Fuchs, 2000a; Marcotti et al., 2003; Tritsch et al., 2007; Johnson et al., 2011). This activity ensures cochlear nucleus neuron survival (Leake et al., 2006), proper wiring of the auditory pathway (Friauf and Lohmann, 1999), and tonotopic maps refinement in the superior olivary nucleus (Kandler, 2004; Clause et al., 2014; Di Guilmi et al., 2019).

During this prehearing period, IHCs receive direct efferent contacts from medial olivocochlear (MOC) neurons, which disappear at hearing onset (Warr and Guinan, 1979; Simmons et al., 1996). This cholinergic input (Glowatzki and Fuchs, 2000; Katz, 2004; Gómez-Casati et al., 2005), mediated by highly calcium-permeable α9α10 nicotinic cholinergic receptors (nAChRs) (Elgoyhen et al., 1994; Weisstaub et al., 2002; Lipovsek et al., 2012), produces inhibition through its coupling to SK2 small-conductance calcium-activated potassium channels (Glowatzki and Fuchs, 2000). Thus, acetylcholine application (Glowatzki and Fuchs, 2000) or electrical stimulation of MOC fibers (Goutman et al., 2005; Wedemeyer et al., 2018) inhibits IHC action potentials (APs) suggesting that efferent inhibition regulates rhythmic activity in the developing auditory pathway (Glowatzki and Fuchs, 2000; Johnson et al., 2011; Sendin et al., 2014; Moglie et al., 2018). However, this idea has been challenged (Tritsch et al., 2010), highlighting that the function of the developmental efferent innervation to the inner ear is still a matter of debate.

Animal models lacking efferent transmission (α9 knock-out, α9KO; Vetter et al., 1999) or exhibiting enhanced efferent activity (α9 knock-in, α9KI; carrying an α9 nAChR mutation that increases suppression of IHC activity; Taranda et al., 2009; Wedemeyer et al., 2018) have been instrumental to a better understanding of the role of the MOC efferent system during development. At the brainstem level, the spontaneous spike patterning of the medial nucleus of the trapezoid body (MNTB) neurons is altered in α9KO mice, leading to a reduced long-term sharpening of functional topography in the lateral superior olive (LSO, Clause et al., 2014). In α9KI mice, the topographic organization of MNTB physiological properties is abolished, and synaptic properties—such as the frequency of miniature and evoked excitatory postsynaptic currents—are altered (Di Guilmi et al., 2019). In addition, MOC modulation influences the correlated activation of inferior colliculus neurons before hearing onset (Babola et al., 2021). Despite these findings, no study has correlated synaptic activity within the same brainstem nucleus as a function of the strength of the MOC efferent modulation. In the present work, we combined electrophysiology with serial block electron microscopy to study the impact of MOC modulation at the same synapse (calyx of Held-MNTB; CH-MNTB) in prehearing mice. Short term synaptic plasticity experiments displayed a significantly higher rate of depression and a smaller ready releasable pool in α9KI mice compared with WT and α9KO. Morphological 3D reconstructions of the CH complexity further demonstrated genotype-dependent differences in CH morphology, including a higher proportion of poly-innervated MNTB cells in α9KI mice.

These findings indicate that enhancing MOC strength drives more pronounced developmental changes synaptic and structural changes in brainstem auditory circuitry than the absence of MOC modulation. Thus, by linking efferent strength to coordinated synaptic and ultrastructural changes at a defined brainstem synapse, this study provides new insight into how MOC activity sculpts developing auditory circuits.

## Materials and Methods

### Animals and experiments

α9KO and α9KI mice has been previously described (α9KO: Vetter et al., 1999; α9KI: Taranda et al., 2009) and were backcrossed with the congenic FVB.129P2-Pde6bþ Tyrc-ch/AntJ strain (https://www.jax.org/strain/004828) for seventeen generations (i.e., N-17). Wild-type (WT), homozygous α9KO and α9KI mice of either sex were used. All experimental protocols were carried out in accordance with the American Veterinary Medical Associations’ AVMA Guidelines on Euthanasia (2013) and approved by the IACUC at INGEBI and the ethical committee of Shanghai Ninth People’s Hospital (No. SH9H-2020-A420-1).

### Electrophysiology on MNTB slices

For slice recordings, 73 mice between 12 and 14 postnatal days (P12-P14) were used. Their brains were removed rapidly after decapitation and placed into an ice-cold low-Ca^2+^ artificial cerebrospinal fluid solution (aCSF). This solution contained the following (in mM): 125 NaCl, 2.5 KCl, 3 MgCl_2_, 0.1 CaCl_2_, 1.25 NaH_2_PO_4_, 0.4 ascorbic acid, 3 myoinositol, 2 pyruvic acid, 25 D-glucose, and 25 NaHCO_3_. The brainstem was glued on a cooled chamber of a vibrating microslicer (Vibratome 1000 Plus, Ted Pella, California, USA). Transverse slices (300 µm thickness) containing the MNTB were sequentially cut and transferred into an incubation chamber containing normal aCSF at 37°C for 30 min. After incubation, slices were allowed to return to room temperature. Normal aCSF had the same composition as the slicing solution except that the MgCl_2_ and CaCl_2_ concentrations were 1 and 2 mM, respectively. The pH was 7.4 when gassed with 95% O_2_ and 5% CO_2_.

### Whole-cell patch-clamp recordings

Slices were transferred to an experimental chamber. During recording, slices were continuously perfused with carbogenated (95% O_2_ and 5% CO_2_) aCSF maintained at room temperature (22–25°C). MNTB neurons were visualized using a Leica DM LFS (Germany) and viewed with differential interference contrast by a 40X water-immersion objective (0.8 numerical aperture water-immersion objective) and a camera with contrast enhancement (DMK 23UP1300, The Imaging Source, North Carolina, USA). Whole-cell recordings were made with patch pipettes pulled from thin-walled borosilicate glass (World Precision Instruments, Florida, USA, RRID:SCR_008593). Electrodes had resistances of 3.8 to 4.5 MΩ. The pipette solution used for isolating synaptic currents was (in mM): 121.3 K-gluconate, 20 KCl, 10 HEPES, 10 phospho-creatine, 0.5 EGTA, 4 Mg-ATP, and 0.3 Li-GTP. The pH was adjusted to 7.3 with KOH. Liquid junction potential was uncompensated in both solutions.

Patch clamp recordings were made using an Axopatch 200B (Molecular Devices, San Jose, CA, USA) amplifier, digitalized by a BCN 2120 board (National Instruments) and acquired with WinWCP software (J. Dempster, University of Strathclyde). Data were sampled at 50 kHz and filtered at 4-6 kHz (low pass Bessel). Series resistances ranged from 6 to 15 MΩ. Whole-cell membrane capacitance (15-25 pF) was registered from the amplifier after compensation of the transient generated by a 10 ms voltage step.

Miniature excitatory post synaptic currents (mEPSCs) were recorded continuously for at least three separate periods of 1 min. Amplitude and frequency were analyzed using Clampfit 10.3 (Molecular Devices, San Jose, CA, USA). Events with a maximal duration of 5 ms and decay time less than 1.5 ms were considered (Taschenberger et al., 2005; Rusu and Borst, 2010). Excitatory post-synaptic currents (EPSCs) were evoked by stimulating (0.1 ms duration and 2–20 mA amplitude) the globular bushy cell axons in the trapezoid body at the midline using a hand-made bipolar platinum electrode and an isolated stimulator (Digitimer DS3; Hertfordshire, UK). Series resistances of EPSC recordings were ∼50-70% compensated. Short term plasticity (STP) was studied by trains of 20 pulses at three different frequencies: 10, 100 and 300 Hz. In each response train, the amplitudes were normalized to the amplitude of the first EPSC. The normalized currents to the first response of each train, was plotted against stimulus number and fitted to an exponential function to obtain both the Plateau (equivalent to the percentage of depression) and Tau (the rate of depression). The release probability (Pr) and the readily releasable pool (RRP) were estimated by the cumulative sum of EPSC amplitudes during the train of 300 Hz and the last ten data points were fitted by linear regression and back-extrapolated to time zero (Schneggenburger et al., 1999). Thus, the zero time intersect gives an estimate of the size of the RRP of synaptic vesicles (N) multiplied by the mean quantal amplitude (q). The release probability can be estimated by dividing the mean amplitude of the first EPSC in the train by the Nq value.

To block Na^+^ currents and avoid postsynaptic action potentials, 10 mM N-(2,6-diethylphenylcarbamoylmethyl)-triethyl-ammonium chloride (QX-314) was added to the internal solution when EPSCs were recorded. Strychnine (1 µM) and bicuculine (20 µM) were added to the aCSF to block inhibitory glycinergic and GABAergic synaptic responses respectively. In experiments in which high-frequency trains were given to study short-term plasticity, the competitive AMPAR antagonist kynurenic acid (1 mM) was added to the aCSF to minimize the impact of postsynaptic receptor desensitization, resulting in an average inhibition of ∼50% of the EPSC amplitude.

### Tissue preparation and staining

We used one animal for each genotype at P12-14 and at P25, with the exception of α9KO at P13 where, due to technical reasons, samples of two animals were collected in two blocks. Animals were anesthetized using isoflurane and perfused transcardially with 0.15 M sodium cacodylate buffer (pH 7.4), followed by a fixation mixture containing 2% paraformaldehyde and 2.5% glutaraldehyde (buffered with 0.08 M sodium cacodylate solution, pH 7.4). An opening was created in the skull by a posterior incision and the brains were kept in the open skull in the fixative solution at 4°C for 48 hours. Then the brainstems were dissected carefully and washed in ice-cold 0.15M cacodylate buffer (pH 7.4). While being submerged in the same buffer, the tissue was cut coronally at a thickness of 400μm using a vibratome (Leica VT 1200S). Slices containing the MNTB were anatomically identified under a dissecting microscopy and collected after careful removal of the surrounding nerve tissues using a razor blade.

Staining of the MNTB samples was performed as described in Hua et al. (2015) and Chequer Charan et al. (2022), with minor modifications. In brief, the samples were washed twice in 0.15 M cacodylate solution (pH 7.4) for 30 min each and sequentially immersed in 2% OsO_4_ (Ted Pella), 2.5% potassium ferrocyanide (Sigma), and 2% OsO_4_ at room temperature (RT) for 1.5, 1.5, and 1.0 hours, respectively. All solutions were buffered with 0.15 M cacodylate solution (pH 7.4). After being washed in cacodylate (0.15 M, pH 7.4) and nanopore-filtered water for 30 min each, the samples were incubated in filtered 1% thiocarbonhydrazide (Sigma) for 1 hour at RT, non-buffered OsO_4_ aqueous solution (2%) for 1.5 hours, 1% uranyl acetate (Sigma) at 50°C for 2 hours, and lead aspartate solution (0.03 M, pH 5.0 adjusted by KOH, EMS) at 50°C for 2 hours, with intermediate wash steps in nanopore-filtered water (twice, 30 min each). Next, the stained tissues were dehydrated through a graded ethanol series (50%, 75%, 90%, 100%, 30 min each at 4°C), transferred from there into pure acetone (three times, 45 min each at RT) and infiltrated with a 1:1 mixture of acetone and Spurr’s resin (4.1 g ERL 4221, 0.95 g DER 736, 5.9 g NSA and 1% DMAE; Sigma-Aldrich) overnight on a rotator at RT. Then the samples were embedded in pure resin for 8 hours and cured in a prewarmed oven (70°C) for 72 hours. The orientation of the MNTB was confirmed based on the morphological landmark delimited by the middle line (medial region) and the VII cranial nerve (lateral region).

### Electron microscopy

The resin-embedded samples were mounted on aluminum metal rivet (Gatan, United Kingdom) and trimmed down (Leica Trim2, Germany) with the MNTB at the center. The block-faces were created by smoothing, using an ultramicrotome (Leica UC7, Germany) equipped with a diamond knife (DiATOME Ltd., Switzerland) and then coated with a thin-layer of carbon using a sputter-coater (Ted Pella, 208C) for quick 2D low-resolution (pixel size: 0.59 µm × 0.59 µm) EM scans (Gemini300, Carl Zeiss; Germany). The orientation and dimension of the MNTB was determined in each sample based on its characteristic soma distribution of large principal neurons. For serial block-face EM imaging, the sample blocks were further trimmed to a smaller block-face (less than 800 μm × 500 μm) and coated with a thin layer of gold (EM ACE600, Leica). Image acquisition of the MNTB was carried out using a commercial SBEM system (Gemini300, Carl Zeiss; 3View2XP, Gatan) in 1-by-4 montage mode with each tile consisting of 7000 × 10000 pixels. Imaging parameters were as follows: incident beam energy 2keV; 15 nm pixel size; pixel dwell time 1.5 μs. Serial sectioning was done at a thickness of 50 nm depending on the sample quality. The in-plane stitching was done in FIJI (plug-in; Preibisch et al., 2009) and consecutive slices were then aligned using a self-written MATLAB script based on cross-correlation maximum according to the dorso-vental (X), medio-lateral (Y) and rostro-caudal (Z) axis. This resulted in the followings EM volumes of (XYZ): 131.5 µm × 153.3 µm × 67.0 µm (WT P14 first block), 132.2 µm × 159.0 µm × 65.5 µm (WT P14 second block), 122.5 µm × 541.8 µm × 92.0 µm (WT P25 first block), 184.4 µm × 208.8 µm × 116.0 µm (WT P25 second block), 157.5 µm × 160.8 µm × 77.0 µm (α9KI P13 first block), 131.3 µm × 158.7 µm × 68.0 µm (α9KI P13 second block), 129.9 µm × 549.6 µm × 107.3 µm (α9KI P25), 128.5 µm × 160.3 µm × 79.3 µm (α9KO1 P12 first block), 123.9 µm × 153.0 µm × 24.1 µm (α9KO1 P12 second block), 125.5 µm × 153.2 µm × 38.1 µm (α9KO1 P12 third block), 126.9 µm × 178.8 µm × 49.5 µm (α9KO2 P12 first block), 129.3 µm × 153.9 µm × 68.0 µm (α9KO2 P12 second block), 138.9 µm × 157.5 µm × 68.2 µm (α9KO1 P21 first block) and 141.3 µm × 160.7 µm × 70.7 µm (α9KO1 P21 second block).

### Manual calyx’s skeleton reconstruction

To create representative skeletons of the CH structure, image stacks in Webknossos (open-source visualization and annotation software; Boergens et al., 2017) were analyzed. These structures were principally identified based on the presence of synaptic boutons or innervations (recognizable by the presence of mitochondria, synaptic vesicles, active zones and axonal cone) and were differentiated from other types based on the presynaptic structure (CH), which has three or more branches (Chequer Charan et al., 2022). The skeletons were built using interconnected nodes following these branches.

### Code accessibility

A custom python routine was made to obtain morphological measures from CHs (available for peer review at the following link: https://zenodo.org/records/18155649?token=eyJhbGciOiJIUzUxMiJ9.eyJpZCI6IjQ0ODQxZTBjLTF mMGYtNDY3OC05YjkyLTI5MGRlYjk4ZWQ2NiIsImRhdGEiOnt9LCJyYW5kb20iOiI3YzkyOTQxZTNjN DcyZDkxYWYxYTJkZDIzNDExOGMzMiJ9.6-KCcCrbHNEdf3X0C27P0KnaTrPQOO-SzDhsHVCEF5CE9HdJhf8QWjeLz-E9xEHd17FPH7MzSZo4lh0CALV, access will be made public upon acceptance of the manuscript). From skeletons created by Webknossos, the NML file which contains the spatial information of the nodes corresponding to one CH was downloaded. The code extracts spatial information of the nodes and connections between them by using Depth-First search (DFS)-based reading, a graph traversal algorithm that explores as far as possible along each branch before backtracking. Based on graph-theory (Chikudo et al., 2023) and applying the Horton approach (Horton, 1945) previously applied in studies of neuronal morphology (Arshadi et al., 2021), the following parameters were calculated as shown in Fig. 3C: number of branching points (nodes located at branch bifurcation), number of ends (last nodes located at the end of every branch), full length (the sum of all the distances between interconnected nodes) and number of branches.

### Calyx complexity classification

Based on a previous complexity classification (Grande and Wang, 2011; Chequer Charan et al., 2022) and due to the fact that a greater calyx length could translate into a larger calyx volume (Spirou et al., 2008), CHs morphotypes were divided into three classes based on the full-length (FL) parameter. We arbitrarily chose three data groups so that the mean error bars did not overlap (see Fig. 4B). Thus, we established FL values less than 260 μm for type 1, 260-340 μm for type 2, and more than 340 μm for type 3.

To validate the statistical robustness of this arbitrary classification scheme, we conducted a post hoc K-means cluster analysis (Grande and Wang, 2011) using Statistics Kingdom website (https://www.statskingdom.com/cluster-analysis.html). This method partitioned all 3D-reconstructed calyces (n = 15) into three clusters based on the full length, the number of branching points, the number of ends and the number of branches. The resulting clusters — designated Group 1, 2, and 3 (see Fig. 4C)— were then compared with the means of the same parameters from our arbitrary types 1, 2, and 3 calyces (see Fig. 4B–C). Each mean value from the arbitrary type classification was then compared with the group clustering to confirm no significant differences for every parameter.

### Calyx and MNTB volume reconstruction

Just for illustrative purposes, five CH-MNTB synapses were selected from the P21 WT segmented dataset to illustrate type 1, type 2 and a type 3 CH (Fig. 4Ai) and mono and poly-innervated morphology (Fig. 7A) as described in (Chequer Charan et al., 2022). Briefly, cured segments corresponding to either CH or a MNTB neuron were downloaded as STL files, then imported as a volumetric reconstruction into Blender 3.2 (Stichting Blender Foundation, Amsterdam. Available at: http://www.blender.org).

### Experimental design and statistical analysis

Experiments were designed to reduce the number of animals and taking into account a balance between the number of samples to accurately perform statistical tests and the ethical guidelines for animal research as described above. Data analyses were done blind to genotype. All statistical tests were carried out with Prism 8 software (GraphPad, La Jolla, CA, USA, RRID: 294 SCR_002798). Prior to performing any analysis, data sets were tested for normal distribution and homoscedasticity. If these assumptions were satisfactorily passed, a parametric test was applied. In these cases, comparisons were made by one-way ANOVA and the statistic “F” value with the associated “p-value” significance was reported. Otherwise, non-parametric Mann-Whitney test was used. Values of p<0.05 were considered significant with the exception of the Bonferroni test (used for correlation test on Table 3), where significance was considered for p values less than 0.00833. Average data were expressed and plot as the mean ± S.E.M. In all cases “n” indicates the number of cells tested and “N” indicates the number of animals. For morphotype proportion, Spearman correlation matrix was performed to detect those statistically correlated parameters with “type” assignment (see Table 3). Kolmogorov-Smirnov test was applied for comparing the cumulative distributions of morphometric parameters (Fig. 3E) and chi square was used to test differences between type proportions and genotypes or poly-innervation and genotypes (Fig. 6). When significant, *ad hoc* Fisher exact test was applied comparing WT vs. α9KI or WT vs. α9KO.

### Drugs and reagents

All drugs and reagents were purchased from Sigma-Aldrich (Saint Louis, Missouri, USA, RRID:SCR_008988) except for TTX which was purchased from Tocris Bioscience (Bristol, UK; RRID: SCR_003689) and QX-314 from Hello Bio (Princeton, USA).

## Results

### Evoked synaptic transmission is reduced in mice with enhanced MOC activity

In a previous study, synaptic alterations on the calyx of Held (CH)-MNTB synapse were evaluated in the α9KI mouse model (Di Guilmi et al., 2019). However, the impact of MOC suppression on this synapse (Fig. 1A) in α9KO mice were not evaluated. To directly compare synaptic properties across mouse models with different strength of the MOC efferent system, both spontaneous (Fig. 1B) and evoked (Fig. 1C) synaptic transmission in WT, α9KI and α9KO mice were analyzed in a new set of experiments. The amplitude and frequency of mEPSCs showed no significant differences in mean amplitude (WT: 50.03± 2.78 pA, n = 14, N = 14; α9KI: 50,22 ± 1.61 pA, n = 14, N = 13; α9KO: 45.57 ± 2.96 pA, n = 13, N = 11; ANOVA, F = 1.86, p = 0.1783; Fig. 1D) or mean frequency (WT: 2.45 ± 0.32 Hz, n = 14; α9KI: 3.13 ± 0.52 Hz, n = 14; α9KO: 2.97 ± 0.56 Hz, n = 13; ANOVA, F = 0.5015, p = 0.6121). Although α9KI and α9KO mice showed greater dispersion (CV_WT_: 0.49, CV_α9KI_: 0.6776, CV_α9KO_: 0.6837) and a trend towards higher frequency, spontaneous transmitter release at the CH-MNTB synapse was not affected by altered MOC function. To analyse MNTB principal neuron EPSCs, responses were evoked by afferent stimulation (Fig. 1E), using a bipolar electrode placed at the midline to evoke an action potential in the presynaptic terminal. Principal neurons of the MNTB receive synaptic input from a single giant calyx terminal that generates the stereotyped calyceal EPSC response, which is independent of stimulus intensity above threshold (Barnes-Davies and Forsythe, 1995). While no significant differences in the EPSC amplitudes were observed between WT (5.08 ± 0.30 nA, n =28, N = 21) and α9KO (4.28 ± 0.32nA, n = 20, N = 16; ANOVA, F = 7.3, p = 0.0013; Tukey test, p = 0.1231), the evoked synaptic currents were lower in the α9KI (3.60 ± 0.25nA, n = 25, N = 21; Tukey test, p = 0.0008; Fig. 1F), indicating that enhanced MOC strength alters evoked synaptic transmission.

**Figure 1.**
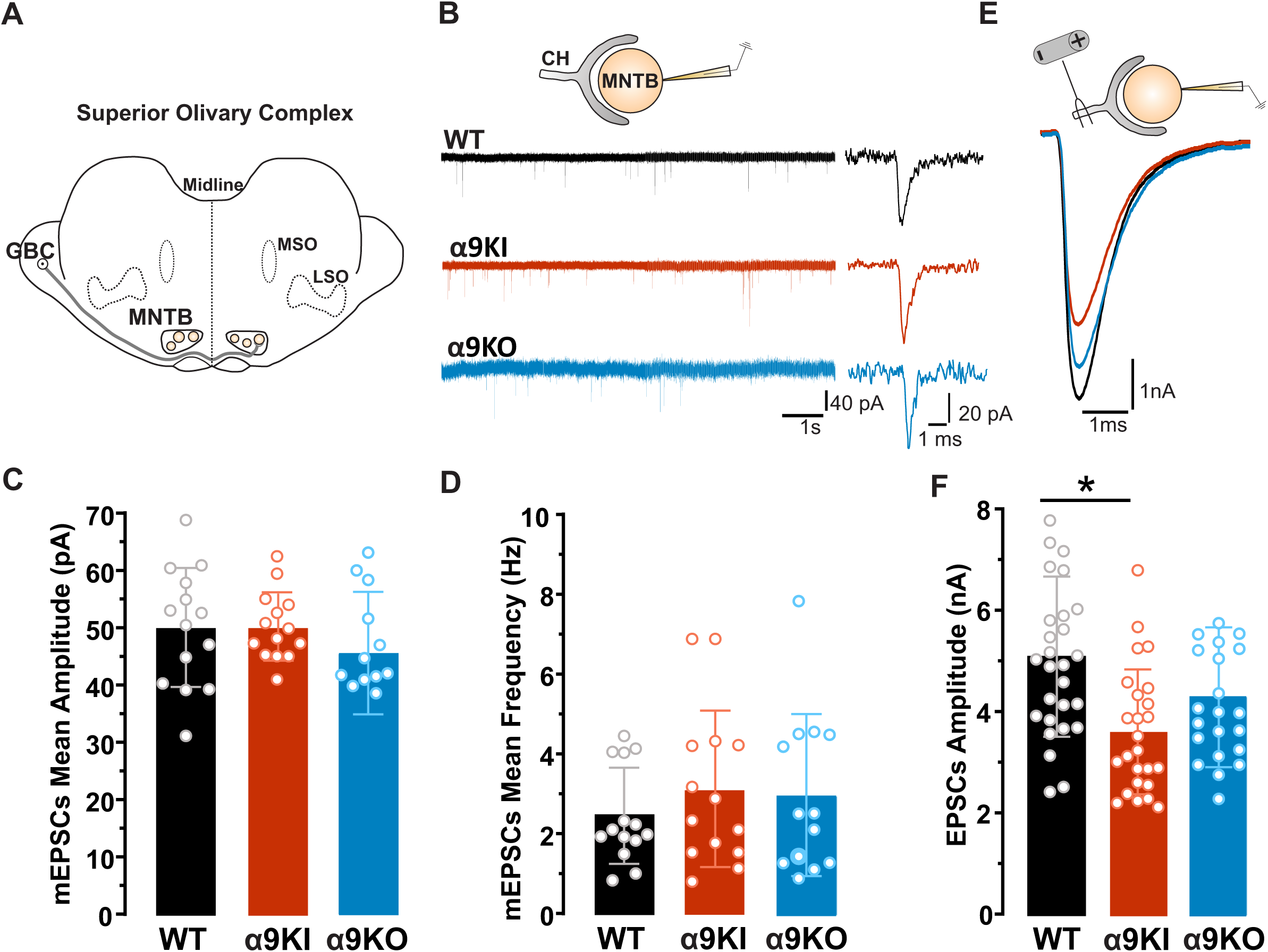
Altered evoked synaptic transmission in α9KI mice. **A.** Schematic diagram of the brainstem at the level of the SOC showing the MNTB localization. **B.** Representative traces of mEPSCs for WT (black), α9KI (red) and α9KO (blue). **C.** The mean mEPSC amplitude for all conditions displayed no significant differences (WT: 50.03± 2.78 pA, n = 14, N = 14; α9KI: 50,22 ± 1.60 pA, n = 14, N = 13; α9KO: 45.57 ± 2.96 pA, n = 13, N = 11; ANOVA, F = 1.86, p = 0.1783). **D.** mEPSC frequency was similar for all genotypes (WT: 2.45 ± 0.32 Hz, n = 14; α9KI: 3.13 ± 0,52 Hz, n = 14; α9KO: 2.97 ± 0.56 Hz, n = 13; ANOVA, F =0.5015, p =0.5015). **E.** Representative traces of evoked EPSCs of MNTB principal cells for WT (black), α9KI (red) and α9KO (blue). **F.** The mean EPSC amplitude was similar in WT (5.08 ± 0.3 nA, n =28, N = 21) and α9KO (4.29 ± 0.32 nA, n = 20, N = 16; ANOVA, F = 7.3, p = 0.0013; Tukey test, p = 0.1231). The mean EPSC amplitude decreased in a9KI (3.60 ± 0.24 nA, n = 25, N = 21; Tukey test, p = 0.0008), when compared to WT.

### Short-term synaptic depression is greater with enhanced MOC activity

To study short term synaptic depression (STD) in the three genotypes, CH were stimulated by trains of either low (10 Hz) or high frequency (100 and 300 Hz) (Fig. 2A, B) to evoke synaptic responses. At 10 Hz, EPSC amplitudes after 2 s of stimulation depressed to 52.28 ± 2.27 % (n = 9, N = 8), 41.93 ± 3.75 % (n = 10, N = 10) and 55.99 ± 4.18 % (n = 11, N = 9) of the first EPSC for WT, α9KI and α9KO mice, respectively (ANOVA, F = 4.135, p = 0.027; Tukey test α9KI vs α9KO, p = 0.0245; Tukey test WT vs α9KO and WT vs Tukey test α9KI, p > 0.05; Fig. 2C). Decay time constants were τ_WT_ = 0.31 ± 0.03 s, τ_α9KI_ = 0.24 ± 0.02 s and τ_α9KO_ = 0.32 ± .02 s (ANOVA, F = 3.628, p = 0.040; Tukey test α9KI vs α9KO, p = 0.05; Tukey test WT vs α9KO and WT vs Tukey test α9KI, p > 0.05; Fig. 2C). During 0.2 s stimulation at 100 Hz, EPSCs from α9KI mice depressed significantly more than those from WT (33.59 ± 1.62%, n = 12, N = 12 for WT and 22.52 ± 2.99%, n = 10, N = 9 for α9KI; ANOVA F = 4.321, p = 0.0224; Tukey test WT-α9KI, p = 0.043) and α9KO (33.92 ± 4.16%, n = 11, N = 8; Tukey test α9KI vs α9KO; p = 0.0381, Tukey test WT vs α9KO, p > 0.05), with similar decay time constants (τ_WT_ = 49.92 ± 3.91 ms, τ_α9KI_ = 33.07 ± 5.92 ms and τ_α9KO_ = 42.51 ± 6.49 ms; ANOVA, F = 2.371, p = 0.1107). This result was also observed at 300 Hz stimulation with larger depression in α9KI compared to WT mice (18.06 ± 1.96%, n = 10, N = 10 for WT and 11.17 ± 1.32%, n = 13, N = 12 for α9KI; ANOVA F=4.607, p = 0.0183; Tukey test WT vs-α9KI, p = 0.0146). No significant difference was found between WT and α9KO (13.21 ± 1.85%, n = 9, N = 6; Tukey test WT-α9KO, Q = 2.75, p = 0.1472). Decay time constants were again similar across groups (τ_WT_ = 15.04 ± 1.91 ms, τ_α9KI_ = 10.57 ± 1.35 ms and τ_α9KO_ = 14.93 ± 1.95ms; ANOVA F = 2.458, p = 0.1032). Taken together, these results indicate that short-term synaptic depression is larger in the mouse model with enhanced strength of MOC inhibition at both low and high frequency stimulation.

**Figure 2.**
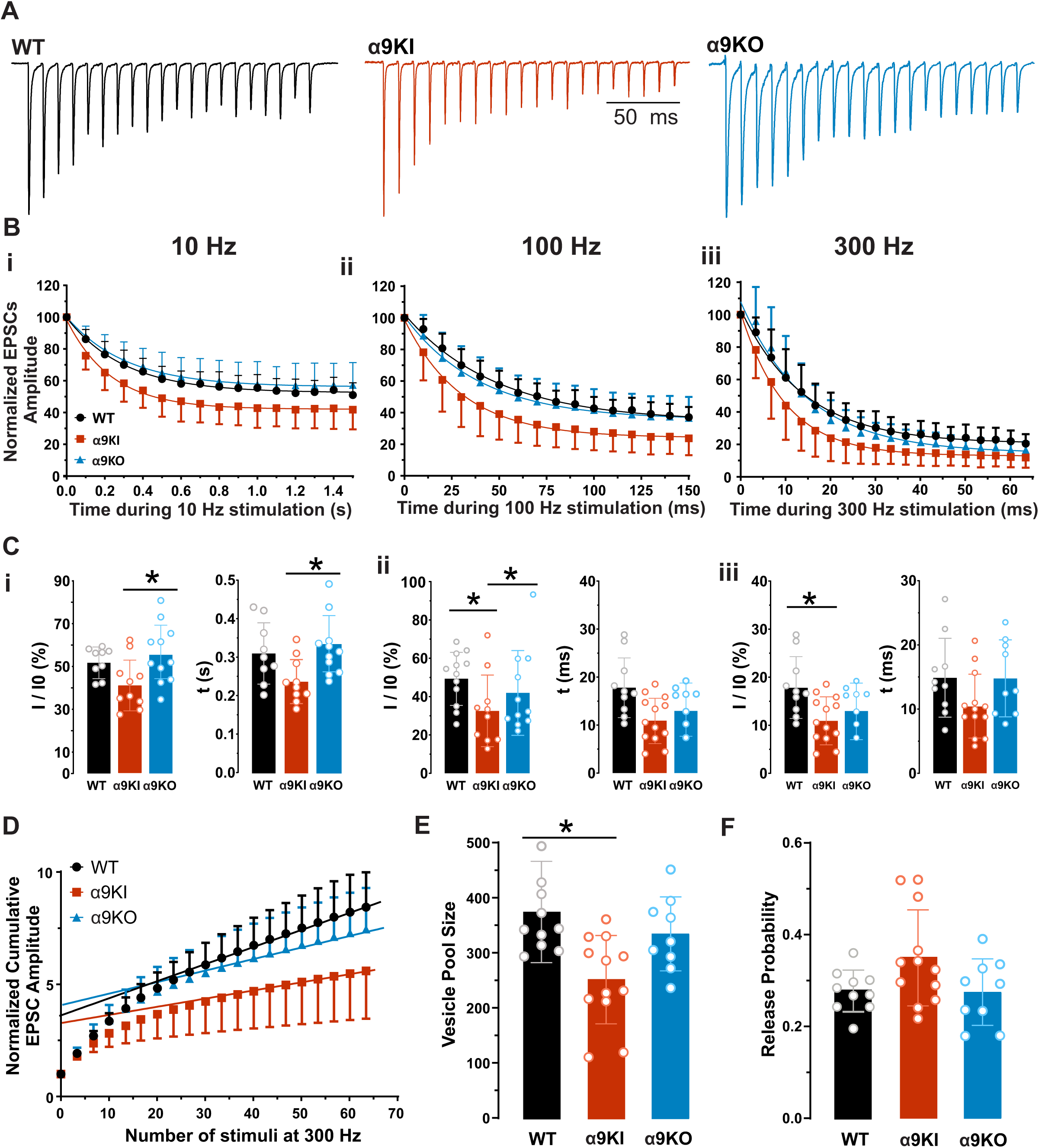
Decreased synaptic strength in the α9KI. **A.** Normalized representative traces showing depression of EPSC amplitudes during 0.2 s stimulation at 100 Hz WT (black), α9KI (red) and α9KO (blue). **B-C/i–iii**, Time course of EPSC depression during either a 10 Hz (**B-C/i**), 100 Hz (**B-C/ii**), or 300 Hz (**B-C/iii**) stimulus train. At 10 Hz, the amplitudes of the EPSCs at the end of the train were 52.28 ± 2.27 % (n = 9, N = 8), 41.93 ± 3.75 % (n = 10, N = 10) and 55.99 ± 4.18 % (n = 11, N = 9) of the first EPSC for WT, α9KI and α9KO, respectively (ANOVA, F = 4.135, p = 0.027; Tukey test α9KI-α9KO, Q = 3.9, p = 0.0245). Decay time constants were τ_WT_ = 0.31 ± 0.03 s, τ_α9KI_ = 0.24 ± 0.02 s and τ_α9KO_ = 0.32 ± .02 s (ANOVA, F = 3.628, p = 0.040; Tukey test α9KI - α9KO, Q = 3.42, p = 0.05). At 100 Hz, the magnitude of depression was larger in α9KI than in both WT (WT: 33.59 ± 1.62%, n = 12, N = 12; α9KI: 22.52 ± 2.99%, n = 10, N = 9; ANOVA F = 4.321, p = 0.0224; Tukey test WT-α9KI, Q = 3.63, p = 0.0403) and α9KO (33.92 ± 4.16%, n = 11, N = 8; Tukey test α9KI - α9KO, Q = 3.74, p = 0.0381). Similar decay time constants were obtained for the three genotypes (τ_WT_ = 49.92 ± 3.91 ms, τ_α9KI_ = 33.07 ± 5.92 ms and τ_α9KI_ = 42.51 ± 6.49 ms; ANOVA, F = 2.371, p = 0.1107). At 300 Hz, the depression was also larger in the α9KI (α9KI: 11.17 ± 1.32%, n = 13, N = 12) compared to WT (WT: 18.06 ± 1.96%, n = 10, N = 10, ANOVA F=4.607, p = 0.0183, Tukey test, Q = 4.00, p = 0.0146). No differences were observed between WT and α9KO (13.21 ± 1.85%, n = 9, N = 6; Tukey test WT-α9KO, Q = 2.75, p = 0.1472). The rate of depression was similar across genotypes (τ_WT_ = 15.04 ± 1.91 ms, τ_α9KI_ = 10.57 ± 1.35 ms and τ_α9KI_ = 14.93 ± 1.95ms; ANOVA F = 2.458, p = 0.1032). **D.** Normalized cumulative EPSC amplitudes during a 300 Hz train for WT (black), α9KI (red) and α9KO (blue). Solid lines indicate the fit of the last ten data points. The intersection of these lines with the ordinate provides an estimate for the cumulative EPSC evoked by the RRP (Schneggenburger et al., 1999). **E.** The vesicle number of RRPs, estimated by extrapolating to the y-axis and dividing the y-intercept value by the amplitude of mEPSCs, was significantly reduced in the α9KI (WT: 374.02 ± 29.11, *n* = 10, N = 10; α9KI: 253.01 ± 21.78, n = 12, N = 11; ANOVA F = 6.24, p = 0.0057, Tukey WT vs α9KI Q = 4.71, p = 0.005) but not for α9KO (323.20 ± 21.40, n = 9, N = 7; Tukey WT vs α9KO Q = 1.57, p = 0.541). **F.** The Pr was similar for WT (0.28 ± 0.01), α9KI (0.35 ± 0.03) and α9KO (0.27 ± 0.02); (ANOVA F = 3.35, p = 0.0527).

To further understand the basis of this enhanced depression, we next estimated the readily releasable pool (RRP) size and release probability (Pr). For any presynaptic terminal, the number of vesicles released is given by the product of the number of vesicles available for release and their release probability. Pr can be evaluated by estimating the fraction of the RRP released by a single AP (Schneggenburger et al., 1999). We measured the cumulative sum of EPSC amplitudes during a train of 20 stimuli at 300 Hz in WT, α9KI and α9KO mice. The last ten data points were fitted with a linear regression and back-extrapolated to time zero (Fig. 2D). The y-intercept gives an estimate of the size of the RRP of synaptic vesicles (*N*) multiplied by the mean quantal amplitude (*q*). The P_r_ can be estimated by dividing the mean amplitude of the first EPSC (for each cell) in the train by the *N*_q_ value. RRP size (Fig. 2E) was significantly smaller in α9KI mice (253.10 ± 21.78, n = 12, N = 11) than in WT (374.02 ± 29.11, *n* = 10, N = 10; ANOVA F = 6.24, p = 0.0057, Tukey WT vs α9KI Q = 4.71, p = 0.005). No difference was found between WT and α9KO (323.20 ± 21.40, n = 9, N = 7; Tukey WT vs α9KO Q = 1.57, p = 0.541. Estimated P_r_ values (Fig. 2F) were 0.28 ± 0.01 for WT, 0.35 ± 0.03 for α9KI and 0.27 ± 0.02 for α9KO mice (ANOVA F = 3.35, p = 0.0527). We conclude that the enhanced short-term depression observed in α9KI mice is accompanied by a significantly smaller vesicle pool size, consistent with impaired presynaptic release capacity under strengthened MOC modulation.

### The morphology of the calyx of Held is more strongly affected in α9KI than in α9KO mice

It is well established that the presynaptic ultrastructure and functional properties of the CH undergo marked developmental refinement to produce a fast and reliable excitatory synapse (for review see (Borst and Soria van Hoeve, 2012). Furthermore, synaptic performance under axonal stimulation depends on calyx complexity (Taschenberger et al., 2002; Ford et al., 2009; Grande and Wang, 2011). Since we found significant genotype-dependent differences in STP, we next asked whether these functional differences were associated with CH morphological alterations. A detailed ultrastructural study was performed, analyzing the CH morphology by serial block electron microscopy (SBEM) (Fig. 3A, B), combined with a custom Python pipeline (see Section “Materials and methods”). This technique allowed us to analyze the entire MNTB ultrastructure with a high spatial resolution. For each animal, we chose 15 - 17 randomly distributed calyces along the MNTB to compare several morphological aspects. For this purpose, we applied graph theory (Chikudo et al., 2023) along with the Horton method (Horton, 1945), which has been used in neuronal morphology research (Arshadi et al., 2021). This structural approach (Fig. 3C) allowed the extraction of the following parameters: the full length (FL; the sum of all branch lengths), the number of branching points (BP), the number of total ends (TE) and the number of branches (TB) (Fig. 3D).

**Figure 3.**
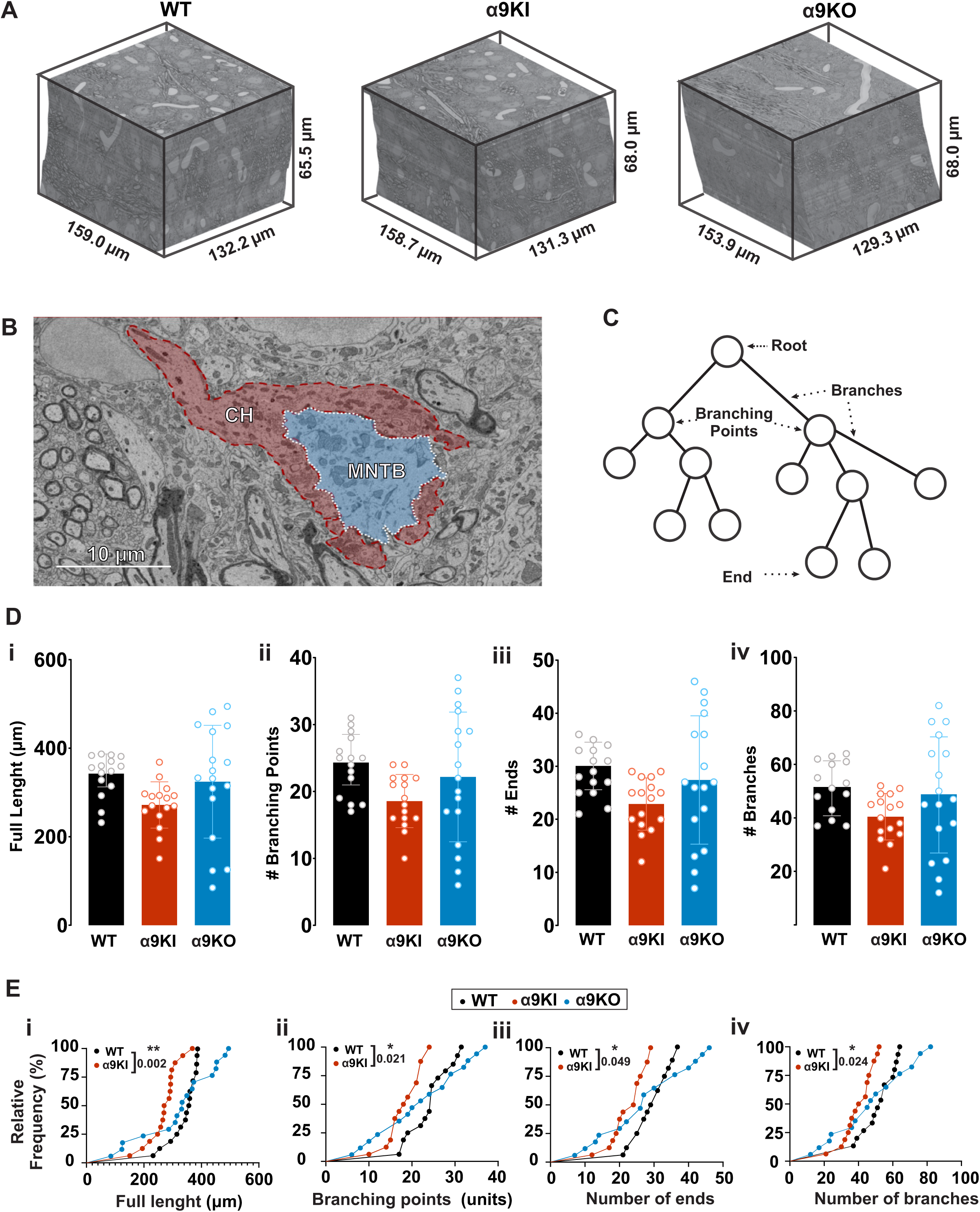
Calyx morphological parameters. **A.** Representative three SBEM volumes of similar dimensions acquired from brainstem coronal slices in WT, α9KI and α9KO. **B.** High-resolution EM image of MNTB principal neurons (blue), which are wrapped by calyx (CH) nerve endings (red). Scale bar 10 μm. Inset: magnification of a representative image showing one calyx terminal onto the cell body of a MNTB neuron. Scale bar 10 μm. **C.** Representative scheme (based on graph theory) indicating the selected parameters for quantifying the complexity of CHs. **D.** Quantification of morphological parameters: full length (FL; **i**), number of branching points (BP; **ii**), number of total ends (TE; **iii**) and number of branches (TB; **iv**). **E.** Cumulative distribution of the FL (**i**), the BP (**ii**), the TE (**iii**) and TB (**iv**) for WT, α9KI and α9KO. Note the right shift in the α9KI towards smaller values. Asterisks denote statistically significant differences, Kolmogorov-Smirnov test (*p < 0.05; **p < 0.005). Error bars indicate the mean ± SEM.

While the mean values of the four parameters did not reach significant differences (Fig. 3D; Table 1), the cumulative distribution of these indexes displayed a clear and significant shift to smaller CHs in α9KI mice compared with WT (Fig. 3E). In contrast, no significant differences were found neither in the mean values (Fig. 3D) nor the cumulative distribution (Fig. 3E) between WT and α9KO, although α9KO values exhibited a notably greater data distribution variability (see CV in Table 1). These results suggest that α9KI calyces are reduced in size, likely reflecting: a) a predominance of simple calyces with few digitations; b) the same proportion of simple and complex structures as the WT but with smaller sizes or c) a combination of both.

**Table 1.** Morphometric quantification of calyces by genotype. Mean value and coefficient of variation (CV) for full length, branching points, total ends and number of branches in WT, α9KI and α9KO.

|  | WT | α9KI | α9KO |
| --- | --- | --- | --- |
| Full lenght - FL [μm] | 339.82 ± 12.32, CV: 0.14 | 271.84 ± 13.07 μm, CV: 0.19 | 324.33 ± 30.88 μm, CV: 0.39 |
| Branching Points - BP (units) | 24.00 ± 1.16, CV: 0.19 | 18.56 ± 0.99, CV: 0.21 | 22.18 ± 2.35, CV: 0.44 |
| Total Ends - TE (units) | 29.13 ± 1.21, CV: 0.16 | 22.88 ± 1.23, CV: 0.21 | 27.41 ± 2.93, CV: 0.44 |
| Number of Branches - TB (units) | 51.80 ± 2.44, CV: 0.18 | 40.44 ± 2.17, CV: 0.21 | 48.59 ± 5.27, CV: 0.45 |
| n | 15 | 16 | 17 |

FL (ANOVA): F=2.708, p=0.077
BP (ANOVA): F=2.336, p=0.1084
TE (ANOVA): F=2.454, p=0.097
TB (ANOVA): F=2.262, p=0.1161

To evaluate these possibilities, a deep ultrastructural study of CH complexity was implemented. CH morphology is heterogeneous and can be clustered into several types according to the gross morphology (Ford et al., 2009) or the number of swellings (Grande and Wang, 2011; Chequer Charan et al., 2022). Such classifications rely on 3D reconstructions using either confocal imaging or segmented electron microscopy data. Here, we took advantage of information extracted from the skeletons and the FL correlates with total calyx volume (Spirou et al., 2008) and used these parameters to generate three groups (see Section “Materials and methods”). We defined FL values less than 260 µm as type 1 (T1), 260-340 µm as type 2 (T2), and greater than 340 µm as type 3 (T3) (Fig. 4A and 4Bi). The remaining morphological parameters (BP, TE and TB) were quantified for each calyx type (Fig. 4Bii-iv and Table 2). We found significant differences and a positive correlation between the type of calyx and each morphometric property (Table 3), indicating that calyx type classification is a good estimator of the morphotype and that the FL works as a good predictor of CH complexity.

**Table 2.** Arbitrary morphometric quantification of different types of calyces by genotype. Mean value for full length, branching points, total ends and number of branches splitting by calyx complexity in WT, α9KI and α9KO.

|  | WT |  |  | α9KI |  |  | α9KO |  |  |
| --- | --- | --- | --- | --- | --- | --- | --- | --- | --- |
|  | Type 1 | Type 2 | Type 3 | Type 1 | Type 2 | Type 3 | Type 1 | Type 2 | Type 3 |
| Full lenght [μm] | 243.33±11.41 | 311.73±10.21 | 367.55±5.05 | 214.13±19.38 | 291.04±6.54 | 368.38 | 132.78±23.52 | 314.86±8.48 | 426.03±19.56 |
| Branching Points (units) | 17.50±0.50 | 20.67±2.19 | 26.30±0.97 | 14.40±1.21 | 20.10±0.86 | 24 | 9.00±1.29 | 19.40±1.29 | 30.50±1.69 |
| Total Ends (units) | 23.00±2.00 | 25.33±2.03 | 31.50±1.04 | 17.20±1.39 | 25.1±0.92 | 29 | 11.00±1.58 | 24.40±1.43 | 37.50±2.44 |
| Total Branches (units) | 37.00±0.00 | 45.00±4.16 | 56.80±1.90 | 30.60±2.60 | 44.20±1.62 | 52 | 19.00±2.79 | 42.80±2.56 | 67.00±4.10 |
| n | 2 | 3 | 10 | 5 | 10 | 1 | 4 | 5 | 8 |

**FL WT (ANOVA):** F=54.14, p=0.0001; Tukey's pT1 vs T2: 0.0016; pT1 vs T3: 0.0001; pT2 vs T3: 0.0006 **BP WT (ANOVA):** F=9.044, p=0.0040; Tukey's pT1 vs T3: 0.0079; pT2 vs T3: 0.00406 **TE WT (ANOVA):** F=8.109, p=0.0059; Tukey's pT1 vs T3: 0.0149; pT2 vs T3: 0.0359 **TB WT (ANOVA):** F=11.6, p=0.0016; Tukey's pT1 vs T3: 0.0028; pT2 vs T3: 0.0276 **FL α9KI (ANOVA):** F=16.96, p=0.0002; Tukey's pT1 vs T2: 0.001; pT1 vs T3: 0.001 **BP α9KI (ANOVA):** F=9.458, p=0.0029; Tukey's pT1 vs T2: 0.0055; pT1 vs T3: 0.0171 **TE α9KI (ANOVA):** F=13.94, p=0.0006; Tukey's pT1 vs T2: 0.0009; pT1 vs T3: 0.0083 **TB α9KI (ANOVA):** F=13.24, p=0.0007; Tukey's pT1 vs T2: 0.0013; pT1 vs T3: 0.0077 **FL α9KO (ANOVA):** F=54.59, p=0.0001; Tukey's pT1 vs T2: 0.0001; pT1 vs T3: <0.0001; pT2 vs T3: <0.0001 **BP α9KO (ANOVA):** F=42.26, p=0.0001; Tukey's pT1 vs T2: 0.0037; pT1 vs T3:< 0.0001; pT2 vs T3: 0.0005 **TE α9KO (ANOVA):** F=33.46, p=0.0001; Tukey's pT1 vs T2: 0.0061; pT1 vs T3: <0.0001; pT2 vs T3: 0.0021 **TB α9KO (ANOVA):** F=38.28, p=0.0001; Tukey's pT1 vs T2: 0.0044; pT1 vs T3: <0.0001; pT2 vs T3: <0.0010

**Figure 4.**
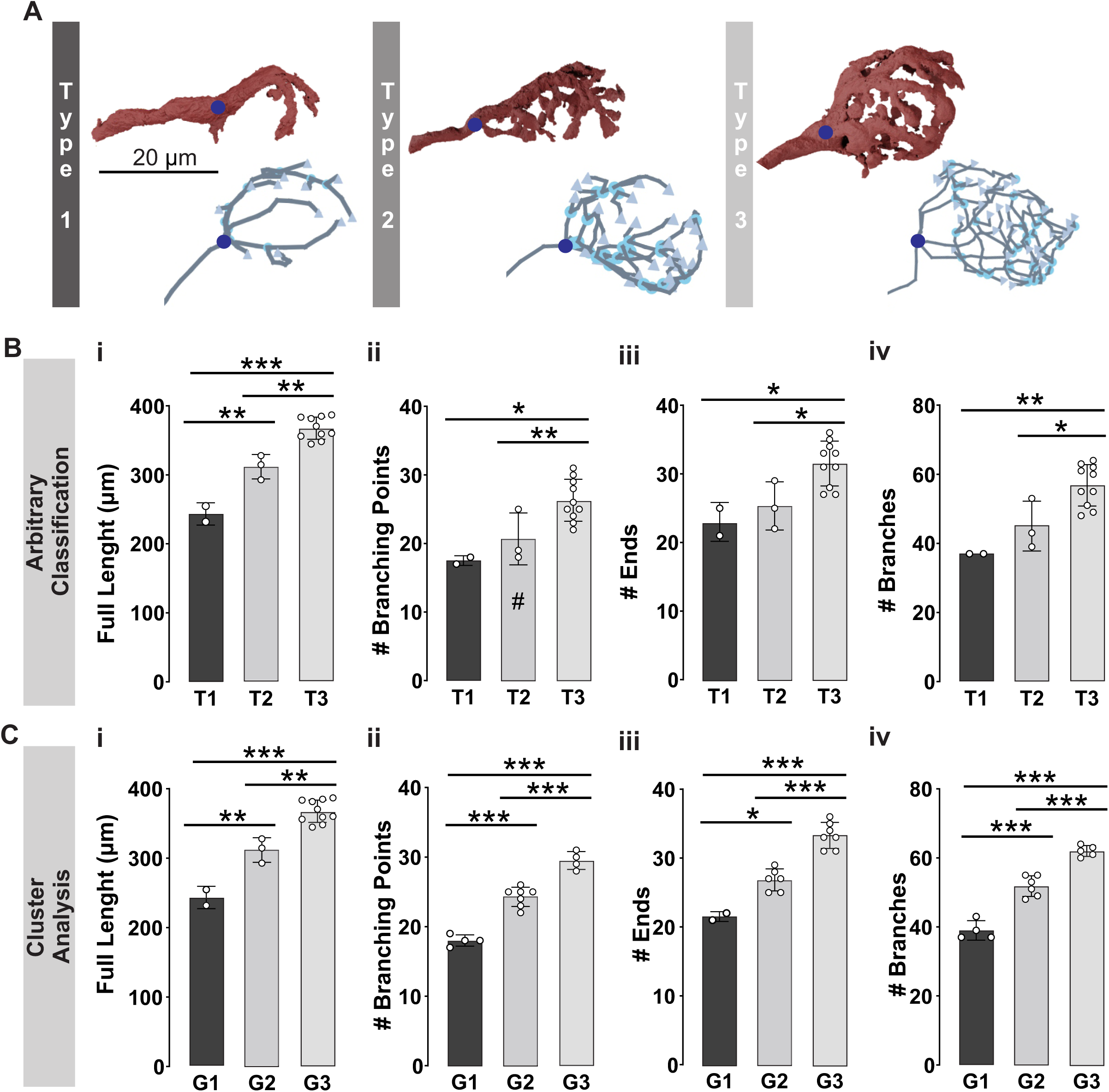
Calyx morphotype classification in WT mice. **A.** Illustrative volume reconstructions showing the heterogeneity in the gross morphology of calyces of Held. Three-dimensional images were reconstructed by Blender Software from segmented images (**upper**). Calyx skeleton reconstruction made with Webknossos used for classification and quantification of different parameters (**lower**). The blue circle corresponds to the axon cone (equivalent to ‘root’ in Fig. 3C), grey triangles the ends and light blue circles the branching points (see also Fig. 3C). Morphological analysis by arbitrary classification (**B**) or Cluster analysis (**C**) of the full length (**i**), the number of branching points (**ii**), the number of total ends (**iii**) and the number of branches (**iv**). Asterisks denote statistically significant differences (Tukey test, \**p* < 0.05; \*\**p* < 0.005, \*\*\**p* < 0.0005). # indicates significant difference between Type 2 (**Bii**) and Group 2 (**Dii**). Error bars indicate the mean ± SEM. For parametric data see Table 2.

To confirm that our arbitrary classification accurately captured the observed morphological heterogeneity, we performed a post hoc cluster analysis (see ‘Materials and Methods’). This analysis grouped the reconstructed CH into three clusters (groups 1 to 3; G1 to G3; Table 4), based on morphological similarity. When partitioning CH by the full length, the branching points, the number of ends and the number of branches, the cluster with the largest values (Group 3; Fig. 4) exhibited significantly larger means than Group 1, mirroring the differences between our arbitrarily defined CH types 1 and 3 (Fig. 4B–C). With exception of the branching points type 2 *versus* group 2 (Tukey post-hoc test, p = 0.047; Fig. 4Bii vs 4Cii), there were no different mean values between the arbitrary classification (types) and the cluster analysis (groups) for simple (T1 versus G1), medium (T2 versus G2) and complex (T3 versus G3) CH. These findings validate that CHs can be classified into distinct morphological phenotypes based on the full length, branching points, number of ends and the number of branches and confirms full length as the best predictor of CH complexity (Spirou et al., 2008).

Once this criterion was established, the same approach was applied for the α9KI and α9KO mice. As observed in the WT, all the parameters selected predicted and positively correlated with the type of complexity both in the α9KI (Fig. 5Ai-iv and Table 2 and 3) and α9KO (Fig. 5Bi-iv and Table 2 and 3), suggesting that our criteria is valid for all genotypes and that morphotype complexity can be explained by the same parameters in the different genetic models.

**Figure 5.**
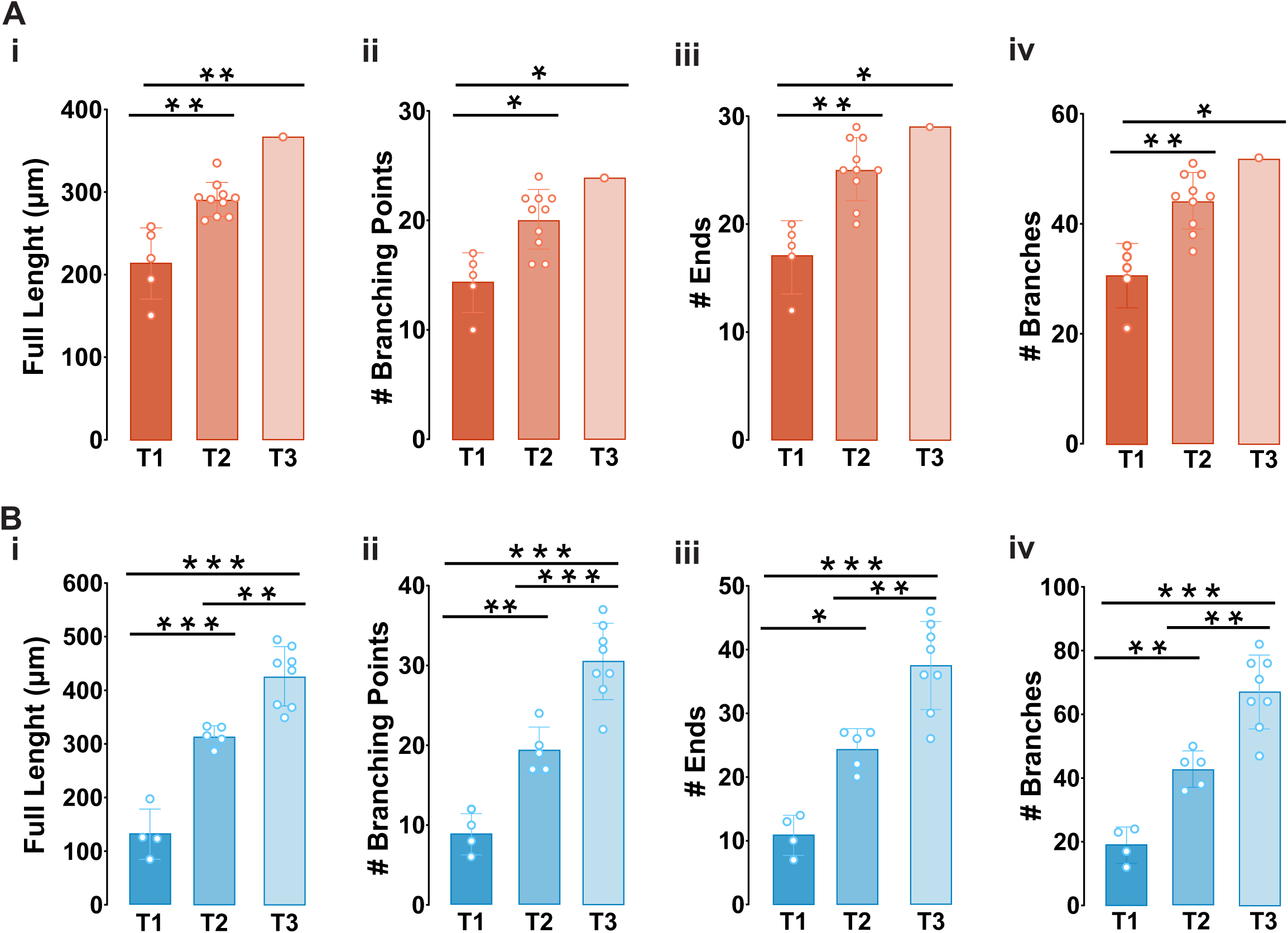
Calyx classification in α9KI and α9KO mice. Morphological analysis of the full length (**i**), the number of branching points (**ii**), the number of total ends (**iii**) and the number of branches (**iv**) for α9KI **(A) and** α9KO **(B)**. Asterisks denote statistically significant differences (Tukey test, \**p* < 0.05; \*\**p* < 0.005, \*\*\**p* < 0.0005). Error bars indicate the mean ± SEM. See also Table 2.

**Table 3.**
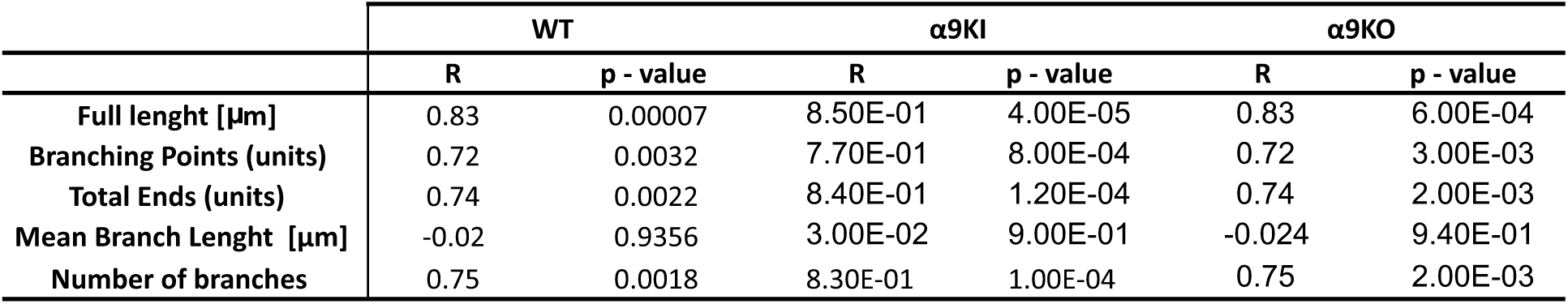
Correlation table comparing several morphometric parameters for each genotype. Spearman correlation coefficients (R) between calyx complexity and morphometric parameters derived from graph theory analyses for each genotype. Due to its negative correlation, the mean branch length parameter was excluded from the classification analysis.

|  | WT |  | α9KI |  | α9KO |  |
| --- | --- | --- | --- | --- | --- | --- |
|  | R | p - value | R | p - value | R | p - value |
| Full lenght [μm] | 0.83 | 0.00007 | 8.50E-01 | 4.00E-05 | 0.83 | 6.00E-04 |
| Branching Points (units) | 0.72 | 0.0032 | 7.70E-01 | 8.00E-04 | 0.72 | 3.00E-03 |
| Total Ends (units) | 0.74 | 0.0022 | 8.40E-01 | 1.20E-04 | 0.74 | 2.00E-03 |
| Mean Branch Lenght [μm] | -0.02 | 0.9356 | 3.00E-02 | 9.00E-01 | -0.024 | 9.40E-01 |
| Number of branches | 0.75 | 0.0018 | 8.30E-01 | 1.00E-04 | 0.75 | 2.00E-03 |

**Table 4.**
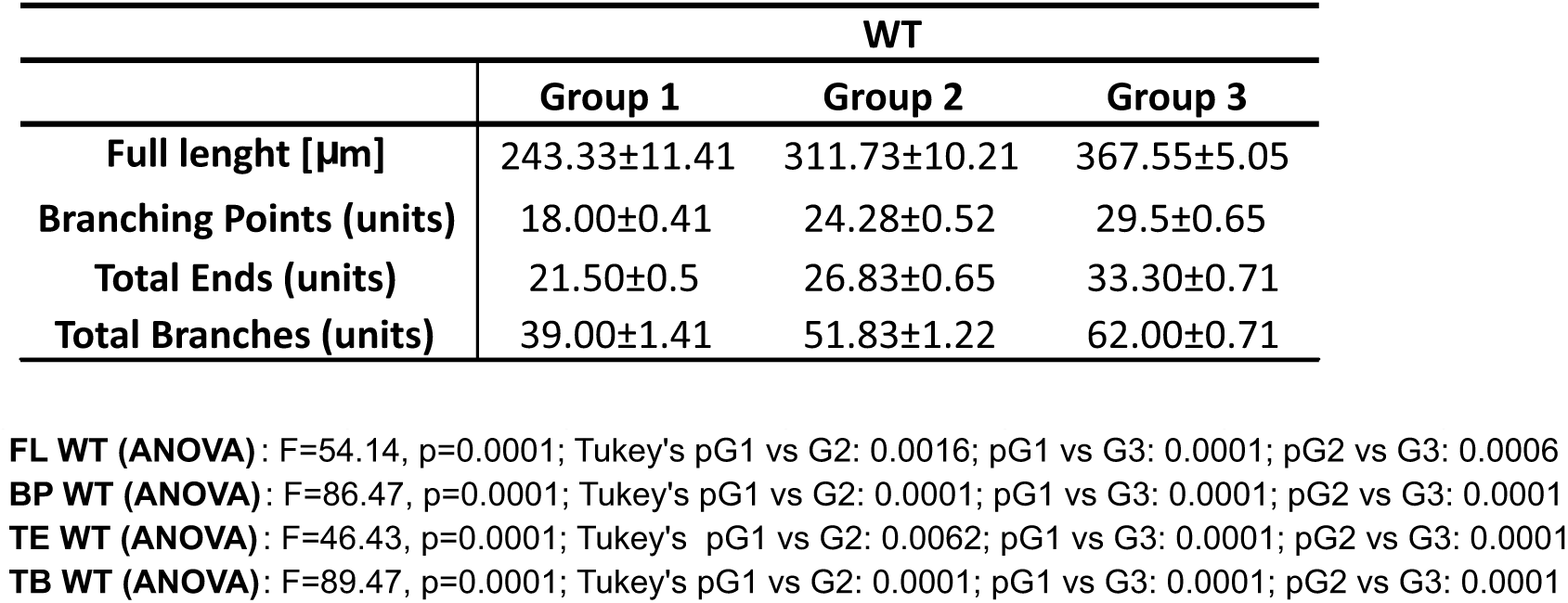
Classification of morphometric parameters by K-mean clustering in WT. Comparison of the full length, branching points, total ends and total branches of calyces assigned to Groups 1, 2, and 3 after a cluster analysis.

We finally compared the percentage of morphotypes by genotype (Fig. 6A) and the pooled type 1/type 2 (Fig. 6B) in order to simplify the complexity comparison. As expected from the previous analysis, where a clear shift towards smaller values was observed (Fig. 3E), a larger proportion of simple CHs (Fig. 6A,B) were detected in α9KI compared to WT (WT _T1-2/T3_: 31 vs 69%; α9KI _T1-2/T3_: 94 vs 6%; χ^2^ = 84.67, df = 1, *p* < 0.0001, ad hoc Fisher test_WT vs α9KI_ : *p* < 0.0001; Fig. 6B). In α9KO, the proportion of simple versus complex types was more balanced (α9KO _T1-2/T3_: 53 vs 47%; ad hoc Fisher test _WT vs α9KO_: *p* = 0,0025), consistent with the wider variability observed in parametric measures (see Fig. 3D,E). Overall, these findings demonstrate that calyx morphotype distribution is strongly affected in the genetically modified mice, highlighting the influence of precise MOC activity during the critical developmental period for achieving a normal morphological maturation of the calyx of Held.

**Figure 6.**
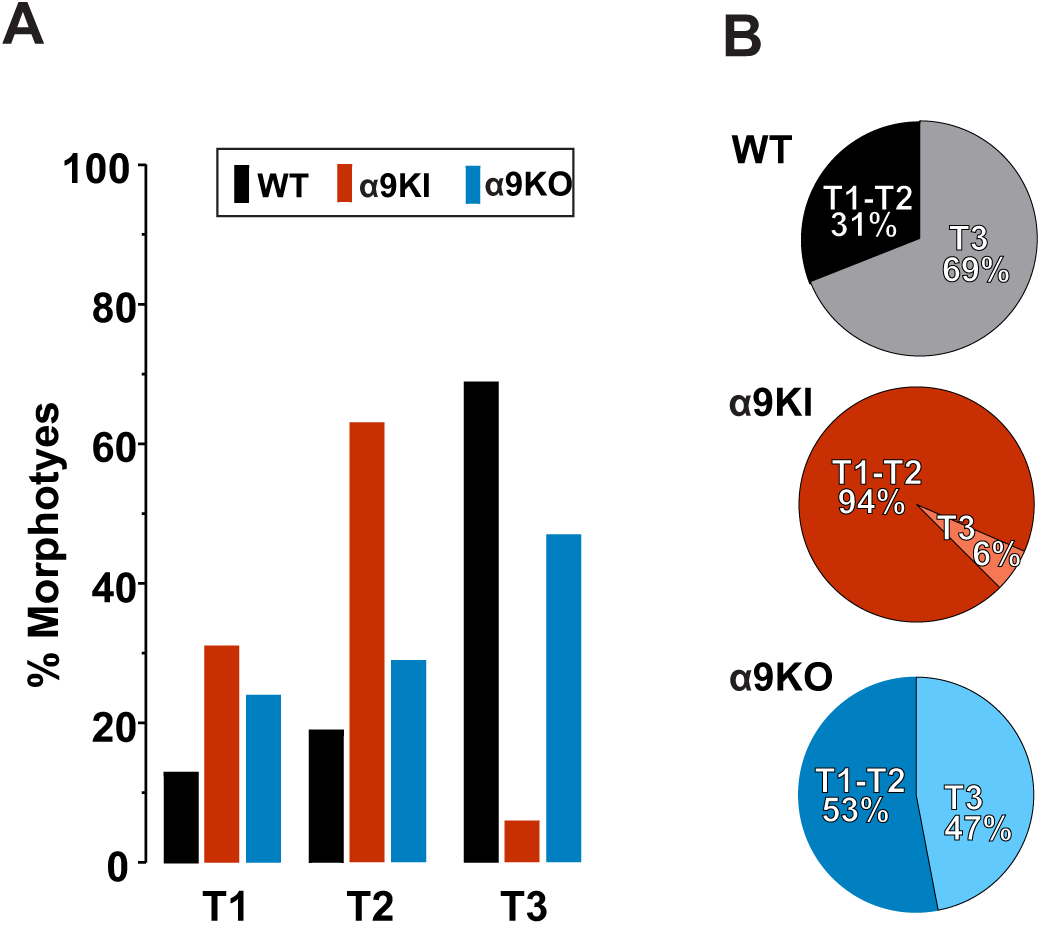
The α9KI mice displayed more simple structures than WT and α9KO. Morphotype distribution across genotypes. **A**. The α9KI and α9KO have more T1 and T2 (T1: 31.25%, T2: 62.50%, T3: 6.25% and T1: 23.53%, T2: 29.41%, T3: 47.06%, respectively) than WT (T1: 12.50%, T2: 18.75%, T3: 68.75%). **B** This behavior is also observed when the morphotype classification is grouped in simple structures (Type 1-2) versus complex ones (type 3)(WT: T1-T2: 31.25%, T3: 68.75%; KI: T1-T2: 93.75%, T3: 6.25%; KO: T1-T2: 52.94%, T3: 47.06%; χ^2^ = 84.74, df = 2, p = 0.0001; Fisher: WT vs. α9KI: p = 0.0001, WT vs α9KO: p = 0.0025.

### Poly-innervated MNTB neurons are more frequent in α9KI mice

During the early postnatal critical period (around the first postnatal week), presynaptic CH terminals make multiple small contacts on MNTB principal neurons, followed by rapid functional and structural refinement (Kandler and Friauf, 1993; Taschenberger et al., 2002; Wimmer et al., 2006; Rodríguez-Contreras et al., 2008). During this pruning process, competing inputs strengthen or weaken until a single dominant calyx remains (Holcomb et al., 2013; Sierksma et al., 2020). In young WT mice (3–4 postnatal weeks), only a subset of MNTB neurons retain multiple calyceal inputs (Milinkeviciute et al., 2019, 2021; Chequer Charan et al., 2022). However, persistent poly-innervation has been reported under pathological or altered developmental conditions, including deletion of bone morphogenetic protein (Xiao et al., 2013), alteration in the microglia (Milinkeviciute et al., 2019, 2021) or enhanced MOC activity (Di Guilmi et al., 2019). In order to analyze poly-innervation we examined the proportion of MNTB neurons receiving more than one calyceal presynaptic terminal at two different ages: P12-P14 and P25 (Figure 7A). At P12-P14 the number of poly-innervated MNTB neurons was approximately 12% of poly-innervated in WT mice [poly-innervated/total number of cells; P12-P14: 3/26 cells (11.54%) Fig. 7B], in agreement with previous reports (Milinkeviciute et al., 2019, 2021; Chequer Charan et al., 2022). However, this proportion was threefold larger in α9KI mice at P12-P14, reaching 32.14% (9/28 cells), consistent with earlier electrophysiological observations where a larger proportion of small-amplitude EPSCs were observed (Di Guilmi et al., 2019). Surprisingly, α9KO mice displayed the similar ratio of poly-innervated cells compared to WT (13.6%; 3/22 cells; χ2 = 15.56, df = 2, p = 0.0004; ad hoc Fisher test, WT vs α9KI: p < 0.0001, WT vs α9KO: p = 0.8339). At P25, α9KI mice still displayed a threefold larger percentage of poly-innervated calyces than WT, whereas α9KO remained similar to WT (WT: 13.3%, 6/45 cells; α9KI: 28.95%, 11/38 cells; α9KO: 8%, 2/25 cells; χ2 = 17.33, df = 2, p = 0,0002; ad hoc Fisher test, WT vs α9KI: p = 0,0087, WT vs α9KO: p = 0,3565). Overall, these observations suggest that synaptic pruning during the critical developmental window is strongly influenced by MOC activity, with enhanced efferent strength (α9KI) producing more pronounced disruptions than the absence of cholinergic regulation (α9KO).

**Figure 7.**
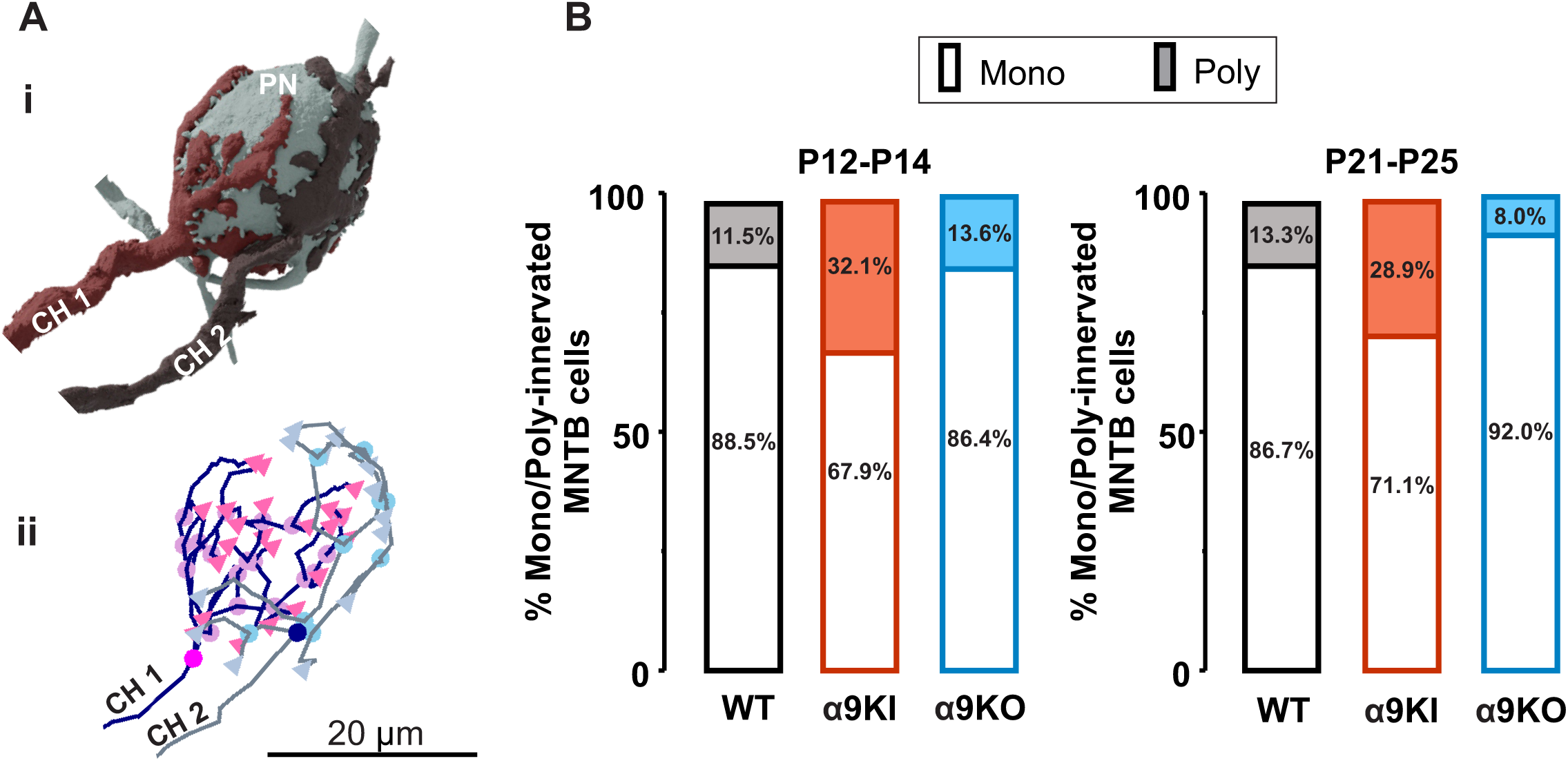
Poly-innervation of MNTB cells is increased in α9KI mice. **(A)** (i) Representative three-dimensional reconstruction of a principal neuron (PN) and its presynaptic calyx of Held (CH) terminal of a poly-innervated (brown and dark gray; CH1 and CH2) MNTB cell and (ii) skeleton reconstruction of presynaptic calyces of Held of another poly-MNTB cell. The pink and blue circle corresponds to the axon cone (equivalent to ‘root’ in Fig. 3C), triangles the ends and light pink and blue circles the branching points (see also Fig. 3C). Scale bar: 20 μm. **(B)** Percentage of poly-innervated cells at different ages. Note that the poly-innervation close to the onset of audition (P12-P14) is near 10% for WT and α9KO, and three-fold higher (∼ 30%) in α9KI mice. This differential proportion is conserved in young adult mice (P21-25).

## Discussion

During early development, the bursting pattern of spontaneous IHC activity is controlled by the MOC system (Johnson et al., 2011; Sendin et al., 2014). Genetic manipulation of α9α10 nAChRs (Vetter et al., 1999, 2007; Taranda et al., 2009) has demonstrated a critical role for MOC efferent input in the maturation of the auditory system. In-vivo recordings from MNTB neurons in α9KO mice revealed that, while the overall bursting activity remained unchanged, the timing of spontaneous spikes is significant altered (Clause et al., 2014), leading to impaired refinement of MNTB-LSO functional maps prior to hearing onset and resulting in deficits in sound localization in adults (Clause et al., 2014). Conversely, genetic enhancement of efferent function (α9KI) disrupts the orderly topographic distribution of biophysical and synaptic properties in the MNTB and causes severe synaptic dysfunction (Di Guilmi et al., 2019). While these studies established the importance of the MOC system for proper central auditory pathway development, none directly compared both the loss (α9KO) and enhancement (α9KI) of MOC activity, combining physiology with morphological ultra-structure, to dissect the changes in the properties of the CH-MNTB synapse in response to developmental altered MOC activity. In the present study we demonstrate that alterations in the strength of the MOC system as seen in α9 nAChR genetically modified mice, produce morphological changes and pronounced synaptic dysfunction in the auditory brainstem. Electrophysiological recordings showed reduced synaptic efficacy in α9KI versus WT (smaller EPSC amplitudes, stronger STD and smaller Pr), but minimal differences in α9KO. Both models showed presynaptic morphological changes, with α9KI exhibiting the greatest deviations: fewer complex CHs and increased poly-innervated MNTB cells. These findings strengthen the view that dysregulation of peripheral MOC activity during the critical period profoundly affects brainstem auditory development (Clause et al., 2014; Di Guilmi et al., 2019).

### Short-term plasticity and morphology

We previously demonstrated that single-evoked EPSCs at the CH-MNTB synapse are altered in the α9KI (Di Guilmi et al., 2019). However, the performance of this synapse during stimulus trains (closely resembling its physiological firing pattern; Sommer et al., 1993; Kopp-Scheinpflug et al., 2003; Hermann et al., 2007; Sonntag et al., 2009) had not been evaluated. Here, we show that STD is larger in α9KI mice at all tested frequencies, consistent with a smaller vesicle pool size, indicating a more profound phenotype in α9KI compared to α9KO mice at the level of the CH. Although detailed spontaneous firing pattern of MNTB neurons in α9KI mice in-vivo remains unknown, global spontaneous activity in the MNTB is reduced (Di Guilmi et al., 2019), consistent with the enhanced cochlear efferent inhibition and reduced cochlear activity (Wedemeyer et al., 2018). Similarly, the lack of afferent responses —caused by knocking down the calcium channel needed for glutamate release by IHCs— leads to synaptic deficits at the CH during development (Erazo-Fischer et al., 2007). Conversely, the suppression of efferent control of IHC (α9KO) maintains the overall frequency of MNTB spontaneous activity unchanged, with a subtle modification in the temporal structure of this activity, crucial for refining the LSO tonotopic map (Clause et al., 2014). The origin of these changes is most likely due to changes in the cochlea where the α9α10 nACHR is expressed and where pre-hearing spontaneous activity is generated. In this regard, Castagna et al., (2025) have demonstrated that both enhanced or absent MOC inhibition disrupts IHC ribbon synapse development, reducing synapse number and altering morphology, suggesting that MOC input plays a key role in shaping synaptic density and preserving pre- and postsynaptic structure. Although ribbon synapse number and morphology differ between WT, α9KO and α9KI, the spatial distribution of synapses along the cochlear axis is preserved in both WT and α9KO but is disrupted in α9KI mice (Boero et al., 2018; Castagna et al., 2025), again indicating a more profound phenotype when suppressing IHC spontaneous activity.

Like most synapses, CH-MNTB synapses undergo extensive structural remodeling during development, involving both pre- and postsynaptic changes (Borst and Soria van Hoeve, 2012). At the pre-synaptic level, the CH structure develops from a single cup-shaped terminal into a fenestrated and more complex structure over the first two postnatal weeks (Kandler and Friauf, 1995; Rowland et al., 2000; Wimmer et al., 2006). Neuronal cytoarchitecture is highly relevant to cellular physiology and, particularly in the MNTB, CH morphology is a major determinant of transmission reliability (Grande and Wang, 2011). MNTB neurons innervated by simple CHs display strong STD, a larger RRP and smaller Pr, whereas those innervated by complex CHs show the opposite profile. Independently of the developmental switch in postsynaptic glutamate receptors (Taschenberger and von Gersdorff, 2000), reduced depression in complex calyces arises (in part) from fenestration, which facilitates efficient glutamate clearance from the synaptic cleft (Renden et al., 2005), representing a morphological adaptation required to sustain the high-frequency synaptic activity of the mature calyx (Sonntag et al., 2009; Crins et al., 2011). Assuming that complex structures have more stalks (branches and digits), greater lengths (Spirou et al., 2008) and consequently more swellings (bouton-like varicosities), where increased calcium accumulation can enhance calcium-dependent synaptic vesicle replenishment (Fekete et al., 2019), it is expected that a larger RRP size would be present in complex CHs (Wang and Kaczmarek, 1998; Wimmer et al., 2006; Fekete et al., 2019). The predominance of simple calyx morphotypes in the α9KI (94%) compared to WT mice (31%), provides a clear explanation for the enhanced synaptic depression and reduced ready releasable vesicle pool. Surprisingly, α9KO—despite altered morphotype proportions—exhibits WT-like depression, suggesting a compensatory physiological mechanism.

### Poly-innervation

Early in development, MNTB neurons receive multiple proto-calyceal inputs from globular bushy cells, which are subsequently pruned to form a single dominant CH by hearing onset (Kuwabara et al., 1991; Hoffpauir et al., 2006; Holcomb et al., 2013; Sierksma et al., 2020). Although in healthy young WT mice (3–4 postnatal weeks), a small subset of MNTB neurons retain multiple calyceal inputs (Milinkeviciute et al., 2019, 2021; Chequer Charan et al., 2022), the persistence of multiple calyceal terminals on a single MNTB neuron beyond the critical developmental window is a hallmark of pathology (Xiao et al., 2013; Milinkeviciute et al., 2019, 2021). Previously, we showed that α9KI mice exhibit a higher frequency of ‘small EPSCs’ compared to WT, a signature of multiple calyceal inputs (Di Guilmi et al., 2019). The present morphological findings provide ultrastructural evidence supporting these electrophysiological findings, since poly-innervation is more prevalent in α9KI compared to WT, with no difference between α9KO and WT.

Unlike the classical model of synaptic competition, in which stronger synapses exhibit a competitive advantage over weaker ones (Cancedda and Poo, 2009), the CH-MNTB synapse is thought to follow a synaptic domination model (Sierksma et al., 2020), where a weak input persists despite coexistence with a dominant synapse. Furthermore, the strongest input already reaches this functional strength while its coverage is still low, suggesting that the increase in strength precedes and instructs the morphological expansion (Hoffpauir et al., 2010; Sierksma et al., 2020). Under this framework, reduced synaptic strength in α9KI should lead to reduced morphological complexity, consistent with our findings. We further propose that MOC-activity contributes to the timing and extent of synaptic domination. Increased poly-innervation as seen in the α9KI (Fig. 7) may lead to a delay in presynaptic domination and pruning. In addition, the persistence of this poli-innervation at least until P25 as seen in the present work, suggests that synaptic dominance must occur during a specific time window and is not compensated after the end of the critical period. Our in depth, detailed morphological ultrastructural study of the CH, reveals cytoarchitecture alterations in the α9KO mice not previously observed (Clause et al., 2014).

### Concluding remarks

The mutation introduced in the α9 subunit to generate the α9KI mouse leads to α9α10 increased channel open probability and to spontaneous channel openings, even in the absence of neurotransmitter and reduced channel desensitization kinetics (Plazas et al., 2005). This is translated into a complete and long-lasting inhibition of spiking activity in α9KI IHCs, compared to the subtle and transient inhibition observed in WT (Wedemeyer et al., 2018), suggesting that spontaneous spiking IHC activity is almost abolished in α9KI. Although in-vivo MOC firing rates remain unknown, tightly regulated IHC spontaneous activity is essential for normal auditory development (Clause et al., 2014; Di Guilmi et al., 2019; Babola et al., 2021). The present results indicate that abolishing this spontaneous activity (α9KI) leads to more severe central synaptic alterations than its dysregulation (α9KO). In this context, the α9KI mouse provides a useful model to understand the consequences of the lack of IHC spontaneous activity during the critical period, a condition that, to our knowledge, has not yet been described.

## Author Contributions

DCH: electrophysiological recordings, manual annotation, 3D reconstruction, Phyton script and electrophysiological and morphological analysis; FW: acquisition of electron microscopy datasets; HW: image and data morphological analysis; YQ: WT/α9KI/α9KO sample processing for microscopy; YH: research co-design and funding; WLG and CCDM: data analysis; ABE: research design, research supervision, writing and funding. MEG-C. and CW: research design and manuscript edition. MNDG: research design and supervision, manuscript writing. All authors contributed to the article and approved the submitted version.

## Funding

This work was supported by Agencia Nacional de Promoción Científicas y Técnicas of Argentina (A.B.E. and M.N.D.G), Consejo Nacional de Investigaciones Científicas y Técnicas (grant PIBAA, M.N.D.G), the Scientific Grand Prize of the Fondation Pour l’Auditionthe (A.B.E.), the National Natural Science Foundation of China (82171133 to Y.H.) and Industrial Support Fund of Huangpu District in Shanghai (XK2019011 to YH).

## Competing Interest Statement

The authors declare no competing financial interests.

## Acknowledgments

We thank Juan D. Goutman for critical reading of the manuscript and fruitful discussions and Claudia Gatto for excellent technical assistance.

## Notes

### Competing Interest Statement

The authors have declared no competing interest.

